# Linking phenotypic and genotypic variation: a relaxed phylogenetic approach using the probabilistic programming language Stan

**DOI:** 10.1101/2024.01.23.576950

**Authors:** Patrick Gemmell

## Abstract

PhyloG2P methods link genotype and phenotype by integrating evidence from across a phylogeny. I introduce a Bayesian approach to jointly modelling a continuous trait and a multiple sequence alignment, given a background tree and substitution rate matrix. The aim is to ask whether faster sequence evolution is linked to faster phenotypic evolution. Per-branch substitution rate multipliers (for the alignment) are linked to per-branch variance rates of a Brownian diffusion process (for the trait) via the flexible logistic function. The Brownian diffusion process can evolve on the same tree used to describe the alignment, or on a second tree, for example a tree with branch lengths in units of time. Simulation studies suggest the model can be well estimated using relatively short alignments and reasonably sized trees. An application of the model in both its one-tree and two-tree variants is provided as an example. Notably, the method is implemented concisely using the general-purpose probabilistic programming language Stan.

## Introduction

The term ‘PhyloG2P’ was coined in a review article [1] summarizing methodologies that link genotype and phenotype by making comparisons across a tree. PhyloG2P methods treat genetic observations (e.g. accelerated molecular substitution rate) in the appropriate context (e.g. lineages where a trait is lost) as evidence in favour of a biological association between genotypic and phenotypic evolution. Such methods have tended to focus on binary traits, although some methods have been developed for continuous traits too [2, 3, 4, 5].

Here I introduce Halcyon, a new statistically motivated PhlyoG2P method for jointly modelling a continuous trait and a corresponding focal multiple sequence alignment. Like the existing method PhyloAcc-C [5], the Halcyon model makes use of a null/background species tree and substitution rate multipliers, but unlike PhyloAcc-C, these substitution rate multipliers can scale the rate of molecular evolution in an arbitrary way on a per-branch basis and could therefore be described as ‘relaxed’ (see [6] for a similar use of the term). The conceit of the Halcyon model is that these substitution rate multipliers can then be linked to the rate of trait evolution using variance rates that are a function of the aforementioned substitution rate multipliers. A systematic coincidence of deviation from a null/background model of sequence evolution and a null/background model of trait evolution may then indicate an interesting biological link between genotype and phenotype.

A notable feature of the implementation of Halcyon is that it makes use of the probabilistic programming language Stan [7], and thus it is both remarkably easy to communicate (Appendix 1) and also benefits from the infrastructure for computational statistics that has been built up by the Stan community. Accordingly, as well as communicating the model, an additional aim of this document is to demonstrate the utility of general-purpose statistical tools for real phylogenetic methods development.

## Methodology

### Model overview

Consider a vector **y** = (*y*_1_, …, *y*_*L*_) of continuous trait measurements (e.g. longevity, beak depth, or systolic blood pressure during pregnancy-induced hypertension) placed at the tips of a rooted tree **T** having *L* leaves, *N* = 2*L −* 1 nodes, and *E* = *N −* 1 edges of length **t** = (*t*_1_, …, *t*_*E*_). Consider also a matrix **X**_*L×S*_ representing a DNA multiple sequence alignment made using genetic observations (e.g. enhancers, promoters, concatenated endogenous retrovirus sequences) from the same *L* species (the rows) at *S* sites (the columns). One would like to mathematically model all this data together in order to describe an association, if any, between the rate of trait evolution and the rate of sequence evolution. The Halcyon model approaches this objective by scaling branch lengths **t** using per-branch substitution rate multipliers **r** = (*r*_1_, …, *r*_*E*_) which are then linked to the rate of change of the trait along any given branch using a functional form *σ*^2^(*r*_*j*_) for the variance rate.

### Sequence evolution

Under the proposed model the evolution of the DNA sequences aligned in **X** is treated in the usual way using a substitution rate matrix **Q**_4*×*4_ so that the probability of transition from nucleotide *a* to *b* on branch *j* of length *t*_*j*_ is expm{**Q** *· r*_*j*_ *· t*_*j*_}_*a,b*_ (see e.g. textbooks [8, 9] for details on the continuous-time discrete-state Markov chain approach to molecular evolution). The tree **T** and matrix **Q** (as well as its associated stationary distribution *π*, used to model unconditional nucleotide frequencies at the root of the tree) are treated as fixed, assumed to have been obtained elsewhere using standard methods such as the `phyloFit` command from the PHAST package [10].

### Trait evolution

Trait evolution is modelled using a distribution whose variance grows with (potentially) scaled branch length. For concreteness, I present the normal distribution, though other distributions might be equally plausible a priori. Under this scheme, a change from trait value *y*_*i*_ to *y*_*j*_ along branch (*i, j*) of length *t*_*j*_ is modelled so that 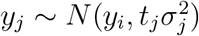,where 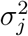 is a trait variance rate obtained using a function somehow dependent on substitution rate multiplier *r*_*j*_. Note that it is possible to use different branch lengths to define the null/background tree used to estimate the substitution rate multipliers and the null/background tree on which the trait evolves. I will refer to a model where two sets of branch lengths are used as a ‘two-tree’ model because even though there is always one tree topology, the two different scaling of the branches will usually look quite different.

It is not immediately clear what function should be used to relate deviations in the rate of sequence evolution (the substitution rate multipliers **r**) to deviations in the rate of trait evolution (the trait variance rates *σ*^***2***^). For this reason, a sensible thing to do is to choose a flexible function, allowing positive, negative, and null associations, as well as various degrees of non-linearity, and the possibility of threshold effects. As Figure 1 shows, the logistic function provides such flexibility. Under this scheme, we have 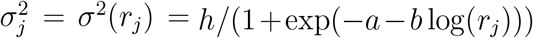 where *h, a*, and *b* are scalar parameters shared across all branches of the tree. When *b* = 0 then the trait is following a Brownian motion on the background phylogeny, as described by e.g. [11]. When *b* ≠0 then the trait is following a Brownian motion that has a variance rate that depends on substitution rate multipliers **r** (that may arbitrarily accelerate or decelerate the rate of nucleotide evolution on any particular branch in order to better describe the alignment **X**) as mediated via the shape of the logistic function.

**Figure 1:**
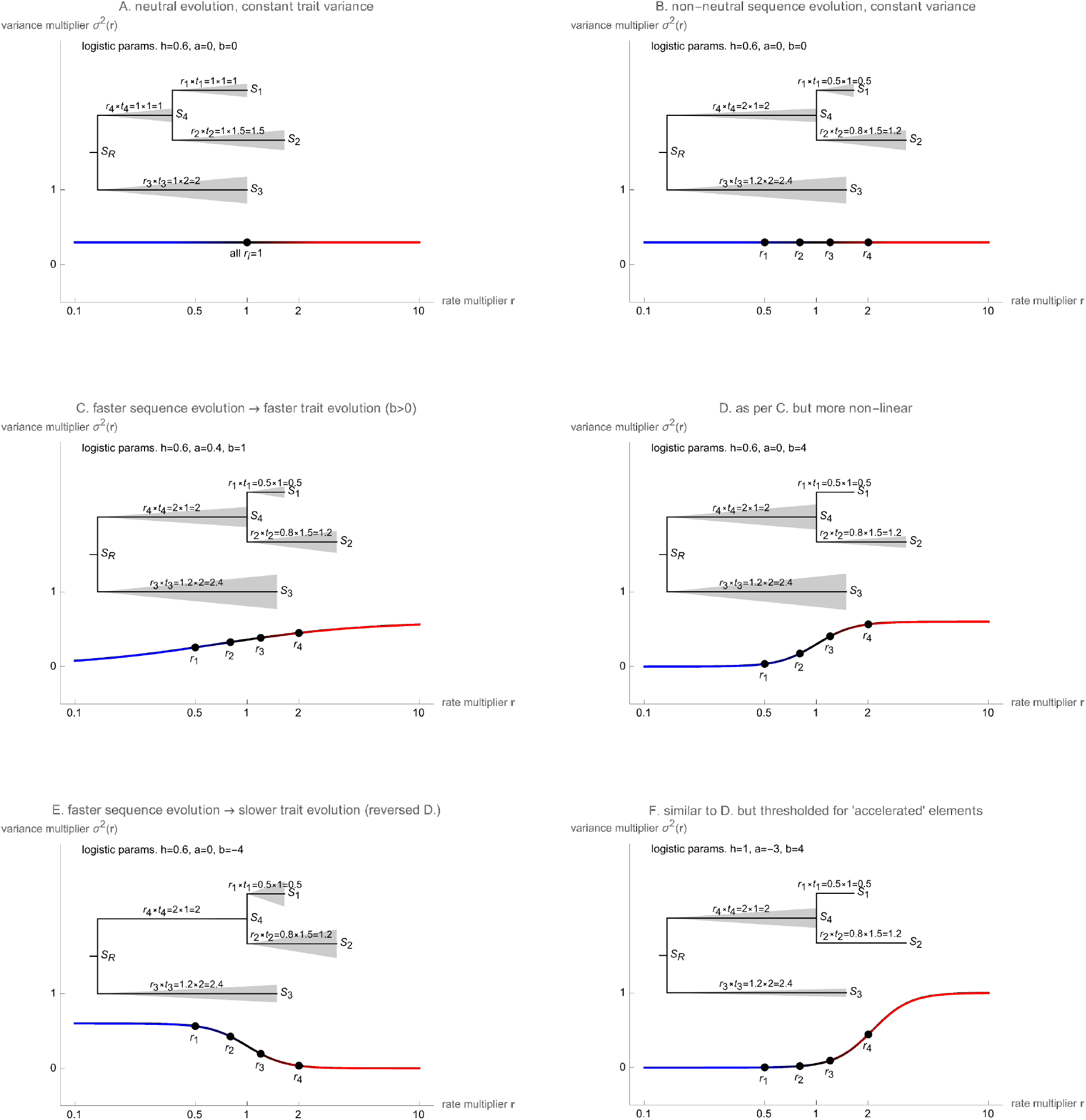
Six example configurations of the Halcyon model based on different parameter values. Outer axes describe variance rate as a function of substitution rate multiplier. Inset in each plot is an example tree, with branch lengths scaled by substitution rate multiplier, and grey triangles showing per-branch trait variance. See the discussion section (below) for an interpretation of the figures assuming the null/background tree **T** is based on the neutral substitution rate. Estimating the model involves considering which parameters are more or less likely based on priors and the observed input data.

### Joint distribution and priors

All together then, using the notational convention that *R* indexes the root node, the joint probability is:

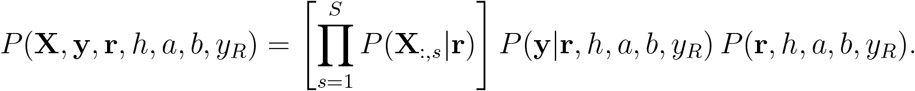

Note that in order to actually calculate a likelihood one marginalizes over all ancestral nucleotides and trait values (Appendix 1).

The following priors were used for simulation studies (below), though one can easily conceive of different choices based on, e.g., the nature of the sequences in alignment **X** or knowledge of *y*_*R*_, the ancestral state of the trait at the root:

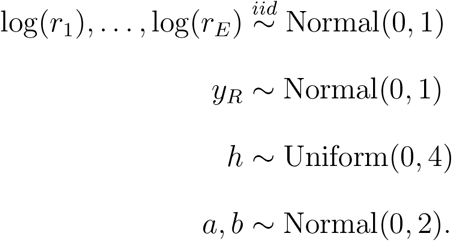

### Implementation

The model was originally investigated using a Metropolis Hastings scheme, a Gibbs sampling scheme, and a Hamiltonian Monte Carlo scheme. The latter approach, implemented using the Stan software, worked well, is described in Appendix 1, and is made available in full at https://github.com/pgemmell/stan_halcyon.

## Results

### Simulation studies

To demonstrate the feasibility of the Halcyon model I undertook six simulation studies (Figure 2) focusing on the problem of estimating parameter *b*, which encodes the presence of a positive or negative association, if any, between the rate of nucleotide and trait evolution. Each simulation study involved 30 replicates. In Simulation A, I drew parameters from their prior distributions and then simulated alignments and trait values under the Halcyon model by making use of functions from the `phangorn::` [12] and `ape::` [13] packages. I used a bifurcating tree with 128 tips, branch lengths of 0.1, and a fairly short alignment length of 120 bp. I then obtained parameter estimates using the `stan_halcyon_logistic.stan` script described in Appendix 1. This involved running 3 randomly initialized MCMC chains per replicate, with random starting values, a warmup of 1,500 samples, and a additional 2,000 samples for parameter estimation. Figure 2 shows one can recover *b* to a reasonable accuracy, and the model appears well calibrated as in 27 of 30 replicates the simulated value was inside the 90% CIs (credible intervals), as is expected. The reported Gelman–Rubin convergence diagnostic for the parameters *h, a*, and *b* was almost always 1.0 (and *<* 1.01 otherwise).

**Figure 2:**
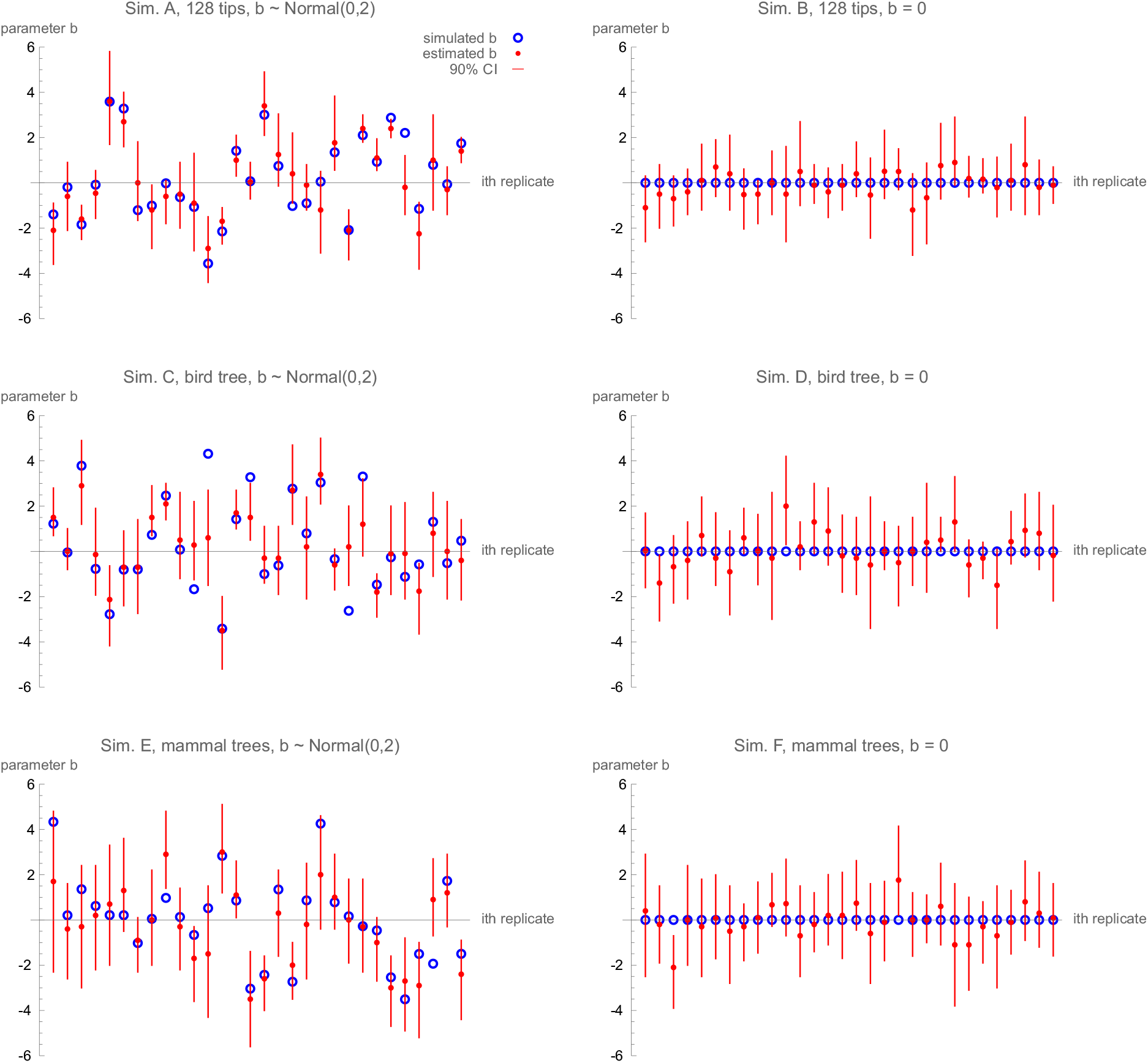
Simulated versus estimated *b* (mean) under model. Simulations were undertaken on a fully bifurcating ultrametric tree with 128 tips and all branch lengths set to 0.1 (row 1), a real-world bird tree with 65 tips (row 2), and paired nucleotide and time trees with 40 tips (row 3). Simulations were performed with an association between the rate of sequence evolution and the rate of trait evolution sampled from the prior (column 1), and with no systematic association (column 2). Simulations used 120 bp alignments.

In Simulation B, with the exception of parameter *b*, I again drew parameters from their prior distributions and then simulated alignments and trait values under the Halcyon model. However, in Simulation B the value of parameter *b* was fixed at 0 to investigate the false positive behaviour of the model. Figure 2B shows that, if anything, the inference was conservative and favoured false negatives, as 30 out of 30 CIs overlap *b* = 0.

Simulations C and D were analogues of Simulations A and B with the difference that a real-world bird tree (Subir Shakya, personal communication) with 65 tips and heterogeneous branch lengths was used in place of the synthetic bifurcating tree with homogeneous branch lengths. Figure 2C shows that estimation of *b* is possible on a smaller, unbalanced tree with irregular branch lengths, and Figure 2D shows that the model still behaves well in the absence of a systematic link between genotype and phenotype. In simulation C, 25 of 30 CIs contained the simulated value of parameter *b*, and in Simulation D 26 of 30 did, again, in both cases, similar to what is expected.

Simulations E and F demonstrate the feasibility of the two-tree concept, the idea that the branch lengths defining the null/background model for the alignment and the branch lengths defining the null/background model of trait evolution need not be the same. In these simulations edge lengths used to describe alignment **X** are expressed in sps (substitutions per site) whereas the variance of the trait under the null/background model is given by a tree with branch lengths in units of time. For these simulations, I downloaded the mammalian tree (originally prepared by Murphy et al. [14]) used to illustrate PhyloAcc-C as well as 10,000 node-dated DNA-only trees from vertlife.org [15]. The latter set of trees have branch lengths in myr (millions of years) and were randomly sub-sampled down to 40 species so that a time tree that matched the topology of the sps tree could be identified. In both simulations E and F, 28 of 30 CIs contained the simulated value of parameter *b*, showing the model remained well calibrated and had a similar false positive rate when used in this new way, even on a yet smaller dataset than used in simulations A–D. In one instance (run 15 of simulation E) a reported Gelman–Rubin convergence diagnostic for parameter *b* was rather large (1.2 not 1.0 as elsewhere), but the estimate of *b* itself was still good.

### Application to a conserved non-coding element

Gemmell et al. [5] identified a conserved non-coding element (labelled VCE277691) where rapid nucleotide evolution was associated with rapid change in the trait long-lived and large-bodied (trait originally prepared by Kowalczyk et al. [16]). Though the Halcyon model is not specific to conserved DNA, it is interesting to see whether a result obtained with a three state model of evolution (PhyloAcc-C) is robust to re-analysis using the more flexible Halcyon model, in either its one-tree or two-tree variant. Examining the two 40 tip trees used in simulations E and F, I observed that on the sps tree the average rate of trait change between two tips connected by exactly two edges varied over about 3 orders of magnitude, from 0.2 per sps (between prairie vole/golden hamster) to 26.6 per sps (human/chimpanzee). Though the time tree had different relative branch lengths to the nucleotide tree, the trait data still varied over a similar range when branch lengths were converted to byr (billions of years), from 0.72 per byr (between prairie vole/golden hamster) or 0.71 (Tasmanian devil/Tammar wallaby) to 31.3 per byr (human/chimpanzee). For this reason, I adjusted the priors on *h, a*, and *b* to the more permissive *a, b ∼* Normal(0, 3) and *h ∼* Gamma(1.5, 0.01) for both analyses. As shown in Figure 3A and B, a strong positive association between rapid sequence evolution and rapid trait evolution was inferred, with an estimated median *b* of 2.0 (90% CI 0.6–3.5) and 1.9 (90% CI 0.4–3.9) under the one and two-tree model, respectively. Nucleotide rate multiplier estimates (Figure 3C and D) were very similar whether the trait was assumed to follow the nucleotide tree or the time tree, and while there were some accelerated branches (e.g. the branch immediately leading to chimp, the branch immediatly leading to pika, rabbit, or their common ancestor, or the branch immediately leading to the great apes), there was no indication that rates fell naturally into a small number of categories; rather the results suggest that there is a great degree of uncertainty over the value of rates on most branches of the tree.

**Figure 3:**
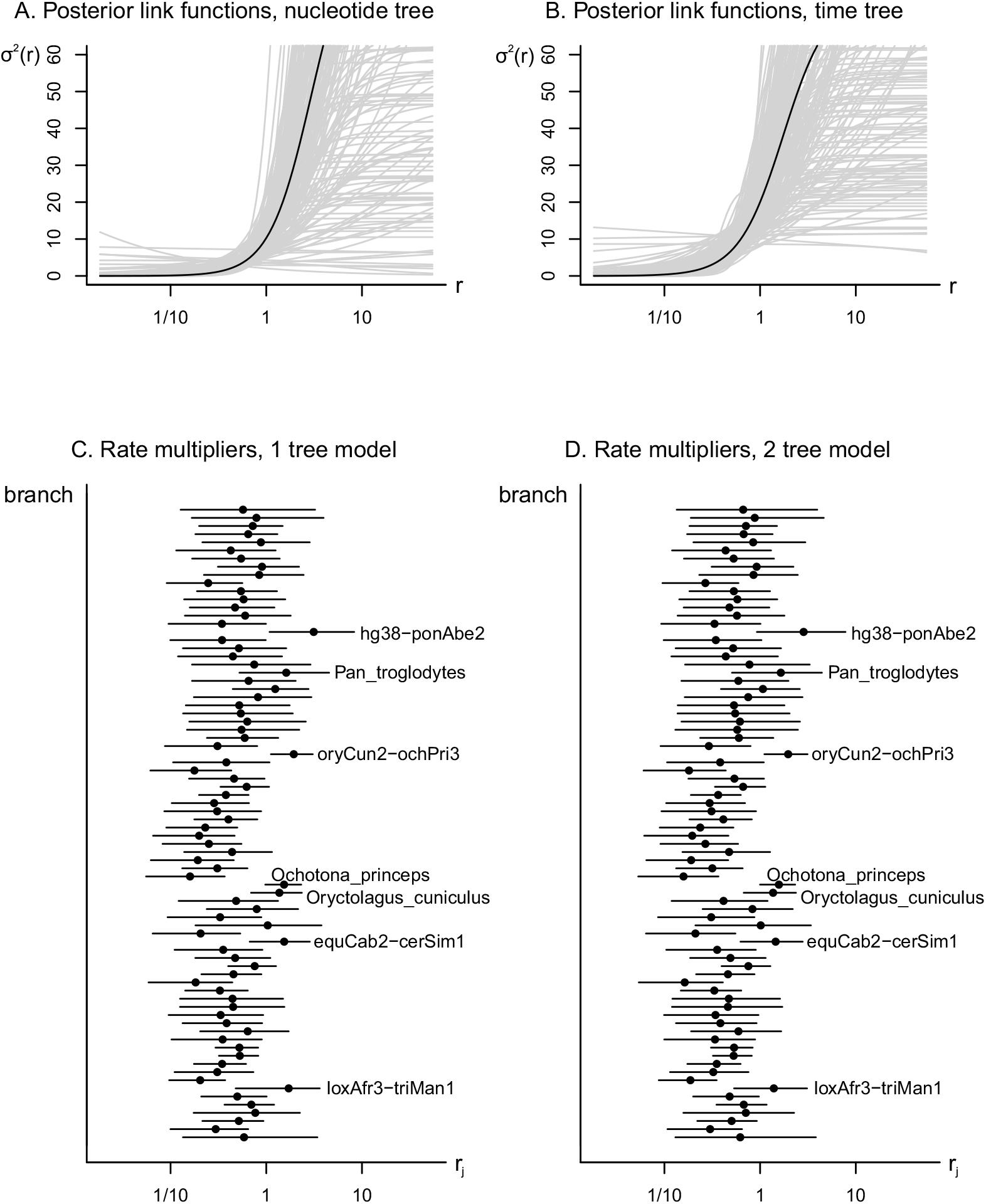
Parameter estimates from inference relating trait long-lived and large-bodied to a conserved non-coding element previously highlighted by Gemmell et al. [5]. Panels A and B summarize parameters *h, a*, and *b* by visualizing the curve they define under the one and two-tree models, respectively. Black lines show curves based on the median inferred values while grey lines show 200 posterior samples. Panels C and D summarize the posterior distribution of rate multipliers under the one and two-tree models, respectively. Black circles represent median values and lines represent 90% CIs.

## Discussion

For presumably practical reasons, popular phylogenetic software has tended to focus on ‘hand-crafted’ Markov chain Monte Carlo samplers (e.g. [2, 17]) rather than make use of general-purpose probabilistic programming languages such as BUGS [18], JAGS [19], and Stan [7]. This is contrary to the widespread use of probabilistic programming languages in other areas of biology (e.g. ecology, biomedical science), as well as in the natural and social sciences (for a diversity of examples see textbooks [20, 21, 22]). While the efficiency of handcrafted samplers can be exceptional, when implementing Halcyon I found the performance of Stan to be ‘good enough’ for challenging phylogenitic inference, which is to say one can get useful work done, as is demonstrated by the inference done above. I was able to reap the benefit of a comprehensive and well tested software tool, with excellent documentation and interfaces (https://mc-stan.org/). Further, a Stan script embodies the notion that ‘programs must be written for people to read, and only incidentally for machines to execute’ (p. xxii) [23], and so is particularly helpful for communicating new models (Appendix 1).

Turning to potential uses of the Halcyon model one wonders how to interpret parameter estimates. The Halcyon model was conceived with the idea of treating *b* as the main parameter of interest. Parameter *b* describes the association between the rate of genotypic evolution and the rate of phenotypic evolution, while parameter *h* captures the ‘height’ of the relationship, and parameter *a* ‘shifts’ the relationship, allowing for threshold effects. Under the model, to the extent the rate of trait evolution is systematically related to a rescaling of the null/background tree via the substitution rate multipliers **r** (needed to explain the alignment **X**) parameter *b* will differ from 0 (e.g. Figure 1C–F). If branches that must be stretched to explain **X** also tend to exhibit faster trait evolution then *b* will tend to be positive (e.g. Figure 1C, D, and F). If branches that must be stretched to explain **X** tend to exhibit slower trait evolution then *b* will tend to be negative (e.g. Figure 1E). This description holds true whether the trait is modelled as evolving on the nucleotide tree used to explain the alignment (a one-tree model, e.g. simulations A–D) or whether it evolves on a separate tree (a two-tree model, e.g. simulations E and F).

The Halcyon model is designed under the assumption that departures from any null expectation of evolution will be sporadic, and of varying magnitude and direction, due to correspondingly unpredictable periods of directional or purifying selection that result from rather random factors. In other words, changes to the same trait in different species will generally be due to different events, that occur at different times, in different places, and unfold in different ways. Therefore correlated rate multipliers are not appropriate even though correlated rates (e.g. [24]) might be an important tool for estimating a null/background tree based on neutral substitution rates.

In the case the null/background tree used to explain the alignment **X** is a genetic tree reflecting an appropriate neutral rate of substitution then rate multipliers **r** themselves do have an interpretation in terms of population genetics, e.g. *r*_*j*_ *>* 1 can be interpreted as directional selection (p. 262–264) [25], as was done by e.g. [26] in the context of retroviral sequence. Note however that the tree used to explain the alignment need not be a genetic tree reflecting the neutral substitution rate, but could instead be a genetic tree representing some other expectation, though in this case the substitution rate multipliers would need to be interpreted differently.

Continuing the discussion under the assumption a tree reflecting neutral substitution rates was used, one could provide the following story about the example configurations in Figure 1. In Figure 1A the focal sequence is evolving neutrally, and there is no systematic link between the rate of genotypic and phenotypic evolution. In Figure 1B the focal sequence may be conserved on some branches (*r*_1_, *r*_2_) and accelerated on others (*r*_3_, *r*_4_), though there is still no systematic link between the rate of genotypic and phenotypic evolution. In Figure 1C parameters are chosen so there is a link between genotypic and phenotypic evolution, and the link is fairly linear. In Figure 1D the relationship is more non-linear than in Figure 1C (there is a saturation effect for both conserved and accelerated elements), while in Figure 1E the relationship is reversed. Figure 1F shows a shifted version of Figure 1D whereby only a substantial acceleration of sequence evolution relative to the neutral rate is linked to an increase in the rate of evolution of a trait. This is in contrast to Figure 1C and D, where an acceleration relative to the most conserved elements is sufficient to induce a change in the rate of phenotypic evolution, as is the case for the model [5]. Note that it is the exploration and weighting of configurations like these example ones that is performed when estimating parameter values of the Halcyon model using Markov chain Monte Carlo.

When the Halcyon model was applied to a conserved non-coding element where the rate of sequence evolution was previously linked to the rate of trait evolution a recapitulation of the positive association between both rates was observed, whether the trait was assumed to evolve on a nucleotide tree (as in the original study) or on a time tree (the two-tree model). In some sense the agreement of the three methods is reassuring: one can argue the identified association is robust to a more flexible model of nucleotide evolution, as well as to the particular species in the tree (the original study used 61, not 40 as used here) and to the null model used. On the other hand, the flexibility is unnerving. There is, as yet, no rule to choose between a nucleotide tree or a time tree as a null/background model for trait evolution. One can argue that the former model is relevant if mutation is a rate limiting step, and that the larger potential phenotypic diversity generated by a bigger population of organisms with shorter generation time is important. On the other hand, if trait evolution tracks a slowly changing environment then perhaps mutations are no longer a limiting factor, and a time tree (as is often used in systematic studies) might be more appropriate. These alternatives suggest that in future the ideas presented in this paper could be developed to address this question by using a model selection framework, or a formulation that interpolates between the two extremes.

How does Halcyon differ from the other Bayesian PhyloG2P methods that model continuous traits? The Halcyon model differs from [5] in that it is much more flexible: it considers arbitrary evolutionary trajectories at the sequence level and therefore also permits more flexible trajectories for traits. It also permits the use of time tree in its two-tree formulation. The Halcyon method differs from the established Coevol method [2] in two main ways. First, Coevol treats the rate of sequence evolution itself as a trait undergoing Brownian motion, and therefore it attempts to auto-correlate rates of evolution across the tree. This makes sense if one expects the rate of evolution of e.g. a mouse and a rat to be more similar to each other than to e.g. an elephant, as the former species are both more genetically related and more physically similar to each other than they are to the latter. In contrast, Halcyon uses rate multipliers. Any similarity in the background/neutral rate of evolution between species can be encoded in the guide tree that is provided as input. Halcyon asks how a focal DNA sequence (i.e. the input alignment) behaves relative to this baseline. Where is the focal sequence accelerated and where is it conserved? The second difference between the two models is how they treat traits. Under both models traits undergo a Brownian motion but under Coevol the question is ‘are fast (slow) rates of nucleotide substitution associated with high trait values?’ Under the Halcyon model the question is ‘are fast (slow) rates of nucleotide evolution associated with high rates of change of a trait?’ The question asked by Coevol seems appropriate for investigating life history traits but the one asked by Halcyon is arguably more relevant for investigating the genetic correlates of evolutionary innovation.

## Acknowledgements

I thank the members of the Jun Liu and Scott Edwards group who asked helpful questions during the Winter 2023 lab meeting. I also thank Timothy Sackton and Subir Shakya for enthusiastic discussions. The computations in this paper were run on the FASRC Cannon cluster supported by the FAS Division of Science Research Computing Group at Harvard University. This work was supported by NIH grant #R01HG011485.

## Appendix

### Pre-processing data for the Stan script

Assume null/background tree **T**, having *E* edges, *L* leaves, *N* = 2*L −* 1 nodes, and a root node index of *R*. Arrange the edges of **T** for post order traversal (e.g. using `ape::postorder`) so that visiting edges in order *E*, …, 1 ensures that ancestral branch (_, *i*) is always visited before its decedents (*i, j*) and (*i, k*).

Construct a boolean matrix **B**_*N×E*_ with **B**_*i,e*_ = 1 if edge *e* is ancestral to node *i* and **B**_*i,e*_ = 0 otherwise. The matrix **B** is useful when calculating a variance-covariance matrix later. For example, use the following (pseudocode) procedure:

~~~
B = zeros(N, E)
for e in E,..,1
  (i,j) = edge[e]
  B[j,:] = B[i,:] // all ancestral edges of i are also ancestral to j
  B[j,e] = 1 // edge e leads directly to node j
end
~~~

The nucleotide alignment **X** can be coded `1–4` for `a, c, g, t`. Trait data may be normalized and centred. The *L* rows of **X** and **B**, and *L* elements of **y** must all share the same order as the tips of the tree.

The edge list, edge lengths, substitution rate matrix **Q**, stationary distribution *π*, and boolean matrix **B** (passed as parameter `path_to_leaf`) are passed to Stan along with the alignment data **X**, and trait data **y**.

### The Stan Halcyon script

Stan models contain a sequence of program statements that define a probability density function conditioned on observed data. These are converted into a sampler using the Stan tool chain.

Stan does not support discrete parameters (e.g. ancestral nucleotides coded as integers) nor directly support phylogenetic inference. However, one can introduce user defined functions using a familiar procedural syntax. For the Halcyon model I introduce two user defined marginalizations and one convenience function.

The first function computes the likelihood of alignment **X** using the Felsenstein algorithm [27], here named `Joe_F()`. The algorithm marginalizes across all possible ancestral nucleotide states. A minor complication to the usual implementation is that under the Halcyon model branch lengths must be rescaled using substitution rate multipliers **r**.

~~~
real Joe_F(array[,] int X, vector r, int L, int N, int E, int R, int S,
  matrix Q, row_vector pi_distro, array[,] int edge, vector edge_length) {
  real log_p_X = 0.0;
  array[S] matrix[N,4] lp_Xup; // prob subtree at i if nuc(i) = j, indexed by site
  for (s in 1:S) {
    for (l in 1:L) {
      for (x in 1:4) {
       if (x == X[l,s]) { lp_Xup[s][l,x] = log(1.0); }
       else             { lp_Xup[s][l,x] = log(0.0); }
     }
   }
   int e = 1;
   while (e <= E) {
    int i = edge[e,1];
    int j = edge[e,2];
    int k = edge[e+1,2];
    real di = r[e] * edge_length[e];
    real dj = r[e+1] * edge_length[e+1];
    matrix[4,4] p_ij = matrix_exp(Q * di);
    matrix[4,4] p_ik = matrix_exp(Q * dj);
    for (x in 1:4) { // x, j, k index rows
       real p_subj = dot_product(p_ij[x], exp(lp_Xup[s][j]));
       real p_subk = dot_product(p_ik[x], exp(lp_Xup[s][k]));
       lp_Xup[s][i,x] = log(p_subj) + log(p_subk);
     }
     e += 2;
   }
   log_p_X += log_sum_exp(lp_Xup[s][R] + log(pi_distro));
 }
 return log_p_X;
}
~~~

The second user defined function, named `sig_fn()`, is a notational convenience that encapsulates the logistic relationship between the variance rates 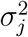 and the substitution rate multipliers **r**. The function is vectorized and returns a variance rate for every branch.

~~~
vector sig_fn(real h, real a, real b, vector r) {
  return h ./ (1 + exp(-a -b * log(r)));
}
~~~

The third user defined function, named `VCV()`, computes the variance-covariance matrix

`Sig` used to calculate the likelihood of seeing trait values **y** at the tips of the tree, conditional on the scaled edge lengths obtained via `sig_fn`. Because `Sig` depends on the values of model parameters *h, a, b*, and **r**, it needs to be constructed repeatedly during model inference.

~~~
matrix VCV(array[,] int Path, int L, int E, vector scaled_edge_length) {
  matrix[L,L] Sig;
  for (j in 1:L) {
    for (k in 1:j) {
      array[E] int common_edges;
      for (ii in 1:E) {
        common_edges[ii] = Path[j,ii] * Path[k,ii];
      }
      real s = 0.0;
      for (ii in 1:E) {
        s = s + common_edges[ii] * scaled_edge_length[ii];
      }
      Sig[j,k] = s;
      Sig[k,j] = s;
      }
  }
  return Sig;
}
~~~

Given the above three functions only one interesting task remains, which is to specify the likelihood of the Halcyon model in a `model` block:

~~~
model {
  h ∼ uniform(0, 4);
  a ∼ normal(0, 2);
  b ∼ normal(0, 2);
  yR ∼ normal(0, 1);
  r ∼ lognormal(0, 1);
  target += Joe_F(X, r, L, N, E, R, S, Q, pi_distro, edge, edge_length);
  matrix[L,L] Sig = VCV(path_to_leaf, L, E, edge_length .* sig_fn(h, a, b, r));
  y ∼ multi_normal(rep_vector(yR, L), Sig);
}
~~~

Modelling the rate of trait evolution using a time tree requires a simple modification of the above model block. Instead of providing one vector named `edge_length` provide two, `edge_length_sps` and `edge_length_byr`. These edge lengths can then be used as follows:

~~~
model {
  …
  target += Joe_F(X, r, L, N, E, R, S, Q, pi_distro, edge, edge_length_sps);
  matrix[L,L] Sig = VCV(path_to_leaf, L, E, edge_length_byr .* sig_fn(h, a, b, r));
  …
}
~~~

